# Anti-microRNA screen uncovers miR-17 family within miR-17~92 cluster as the primary driver of kidney cyst growth

**DOI:** 10.1101/405100

**Authors:** Matanel Yheskel, Ronak Lakhia, Andrea Flaten, Vishal Patel

## Abstract

Autosomal dominant polycystic kidney disease (ADPKD) is the leading genetic cause of renal failure. We have recently shown that inhibiting miR-17~92 is a potential novel therapeutic approach for ADPKD. However, miR-17~92 is a polycistronic cluster that encodes microRNAs (miRNAs) belonging to the miR-17, miR-18, miR-19 and miR-25 families, and the relative pathogenic contribution of these miRNA families to ADPKD progression is unknown. Here we performed an *in vivo* anti-miR screen to identify the miRNA drug targets within the miR-17~92 miRNA cluster. We designed anti-miRs to individually inhibit miR-17, miR-18, miR-19 or miR-25 families in an orthologous ADPKD model. Treatment with anti-miRs against the miR-17 family reduced cyst proliferation, kidney-weight-to-body-weight ratio and cyst index. In contrast, treatment with anti-miRs against the miR-18, 19, or 25 families did not affect cyst growth. Anti-miR-17 treatment recapitulated the gene expression pattern observed after miR-17~92 genetic deletion and was associated with upregulation of mitochondrial metabolism, suppression of the mTOR pathway, induction of autophagy, and inhibition of cyst-associated inflammation. Our results argue against functional cooperation between the various miR-17~92 cluster families in promoting cyst growth, and instead point to miR-17 family is the primary therapeutic target for ADPKD.

## Introduction

Autosomal dominant polycystic kidney disease (ADPKD), caused by mutations in either *PKD1* or *PKD2*, is one of the most common monogenetic disorders and the leading genetic cause of end-stage renal disease (ESRD) in the United States. ^1-3^ This condition is characterized by the presence of numerous fluid-filled cysts originating from renal tubules. Excessive proliferation of mutant epithelial cells causes the cysts to expand eventually leading to impairment of normal kidney function.

microRNAs (miRNAs) are small, 22-nucleotide long, non-coding RNAs that post-transcriptionally inhibit mRNA expression.^4^ Watson-Crick base-pairing between the seed sequence, a stretch of 6 nucleotides (2 through 8) at 5’ end of a mature miRNA, and a complementary sequence on target mRNA results in the repression of that mRNA.^5^ miRNAs that harbor the same seed are classified into one family. miRNAs belonging to the one family have redundant biological functions because they target the same mRNAs. In diseases such as cancer ^6-9^, and as we have recently shown in ADPKD^10, 11^, aberrant activation of miRNAs can drive disease progression. Accordingly, synthetic oligonucleotides known as anti-miRs have emerged as a novel therapeutic platform to inhibit pathogenic miRNAs.^12, 13^ Anti-miRs harbor complementary sequences and Watson-Crick base pair to cognate miRNAs thereby sterically hindering their function.^14^ In general, anti-miRs have a long half-life (>21 days) and are delivered primarily to the liver and kidney making them attractive therapeutic agents to treat chronic diseases such as ADPKD.

We have previously shown that overexpression of the miR-17~92 cluster in normal kidneys is sufficient to produce cysts.^10^ Conversely, expression of the miR-17~92 cluster is increased in human and murine ADPKD, and its deletion reduced cyst burden in four orthologous mouse models of ADPKD.^15^ Based on this work there has been a growing interest in developing anti-miR drugs designed to target the miR-17~92 cluster.^16-21^ miR-17~92, however, is a complex polycistronic cluster that encodes 6 individual miRNAs (miR-17, miR-18a, miR-19a, miR-20a, miR-19b-1, and miR-92a-1). Moreover, the mammalian genome encodes two additional paralogous clusters known as miR-106a~363 and miR-106b~25. The miR-106a~363 encodes 6 miRNAs (miR-106a, miR-18b, miR-20b, miR-19b-2, miR-92a-2, and miR-363) and the miR-106b~25 cluster encodes 3 (miR-106b, miR-93, and miR-25). miR-92b is independently transcribed.^22^ Collectively, these 15 miRNAs encoded from the four loci can be categorized into 4 miRNA families: miR-17, miR-18, miR-19, and miR-25.^23^ The relative pathogenic contribution of each miRNA family to ADPKD progression is not known. Therefore, which miRNA families should be the focus of drug development remains unclear.

To address this question, we used anti-miRs to selectively inhibit the expression of each miRNA family in an orthologous model of ADPKD. We report that only the inhibition of miR-17 family, but not the miR-18, miR-19, or miR-25 families, attenuates cyst growth. Similar to genetic miR-17~92 deletion, we found that inhibition of the miR-17 family improved mitochondrial metabolism and cyst-associated inflammation. In addition, anti-miR-17 treatment provided the additional benefit of reduced mTOR signaling. These data suggest that, within miR-17~92 cluster, the miR-17 family is the primary therapeutic target for ADPKD.

## Results

### Expression of miRNAs produced by the miR-17~92 and related clusters in ADPKD mouse model

We used the KspCre/*Pkd1*^*F/RC*^ (*Pkd1*-KO) mouse model for our studies. *Pkd1*-KO is an orthologous ADPKD model that harbors a germline hypomorphic *Pkd1* mutation (R3277C)^24^ on one allele and *loxP* sites flanking *Pkd1* exons 2 and 4 on the other. We used KspCre-mediated recombination to produce a compound mutant mouse with a kidney-specific null mutation on one allele and a hypomorphic mutation on the other. This is an aggressive, but long-lived model of ADPKD with a median survival of about 6 months.^15^ We began by comprehensively analyzing the expression levels of each mature miRNA encoded by the miR-17~92, miR-106a~363, and miR-106b~25 clusters in kidneys of *Pkd1*-KO mice at 3 weeks of age. Quantitative real-time PCR (Q-PCR) analysis revealed that five of the six mature miRNAs derived from the miR-17~92 cluster were upregulated in cystic kidneys compared to control kidneys **(Fig. 1A)**. miR-17 and miR-18a were the most upregulated showing an increase in expression by 133.2% and 142.9%, respectively. The expression of miR-19a, miR-20a, and miR-19b were increased by 43.6%, 46.7%, and 29.3%, respectively. miR-92a expression was unchanged. All mature miRNAs derived from the miR-106b~93 cluster were also upregulated. Expression of miR-106b was increased by 45.5%, miR-25 by 37.2%, and miR-93 by 29.4% in cystic kidneys compared to control kidneys **(Fig. 1B)**. miR-92b is transcribed independently and its expression was increased by 66.9% **(Fig. 1C)**. miRNAs derived from the miR-106a~363 cluster exhibited more variable expression **(Fig. 1D)**. miR-106a level was increased by 51.2% whereas miR-18b, miR-20b, and miR-363 was decreased by 53.8%, 63.9% and 59.4%, respectively. Thus, five out of six miRNAs belonging to the miR-17 family, one out of two miRNAs belonging the miR-18 family, both miRNAs belonging to the miR-19 family, and two of four miRNAs belonging to the miR-25 family were upregulated in cystic kidneys compared to control kidneys.

**Figure 1:**
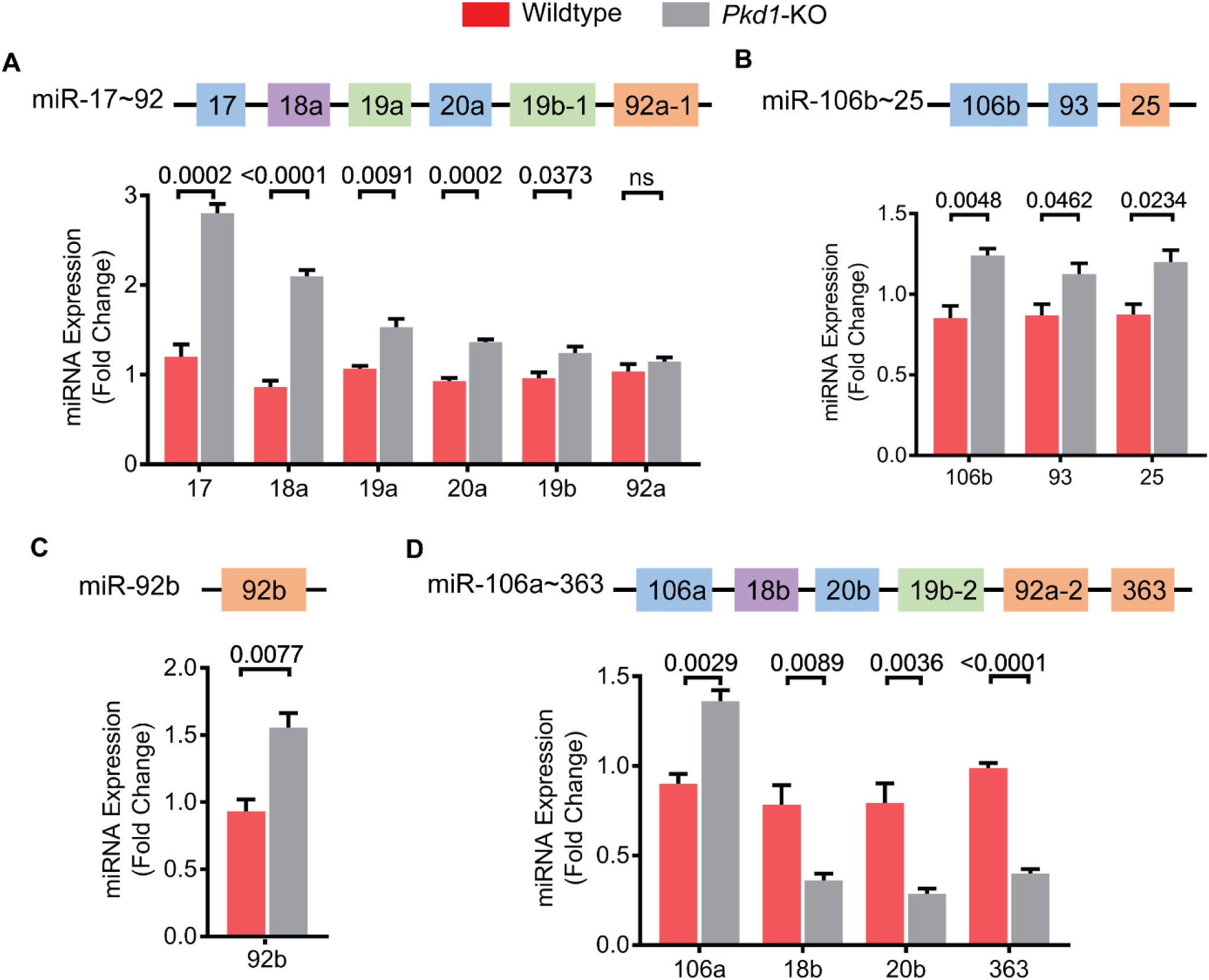
Expression of miR-17~92 and related clusters in *Pkd1*-KO kidneys. The miR-17~92, miR-106a~363, miR-106b~25, and miR-92b encode miRNAs belonging to four families: miR-17 (blue), miR-18 (purple), miR-19 (green), and miR-25 (orange). miRNAs are color-coded based on their family. **(A-C)** Q-PCR analysis showed that the expression of five of the six miRNAs derived from the miR-17~92 cluster, all three miRNAs derived from the miR-106b~25 cluster, and the independently transcribed miR-92b are upregulated in *Pkd1*-KO compared to control kidneys. **(D)** miRNAs derived from the miR-106a~363 cluster showed variable expression. miR-106a expression was increased whereas miR-18b, 20b, and miR-363 expression was decreased. Wildtype n=3, *Pkd1*-KO n=5. All mice were 3 weeks of age. Data are presented as mean ± SEM. Statistical analyses: Student’s t-test, ns indicates *P*>0.05.

### Anti-miRs specifically inhibit cognate miRNA family without affecting the expression of unrelated miRNAs

We used 12 to 16 nucleotides long, locked nucleic acid-modified anti-miRs to selectively inhibit either the miR-17, miR-18, miR-19, or miR-25 family. Anti-miRs were designed to Watson-Crick base pair with the majority but not the entire length of cognate mature miRNA sequence. The nucleotide sequences of anti-miRs used in the current study are shown in Supplementary Table 1. A mixture of five anti-miRs was used to inhibit all six miR-17 family members simultaneously. Each anti-miR in the mix harbored a perfectly complimentary sequence to the seed sequence (shown in bold in Supplementary Table 1) of miR-17 family. However, the flanking nucleotide sequences were slightly modified to account for minor differences in the sequences of the various miR-17 family members. Using a similar strategy, we designed three anti-miRs that collectively targeted all four miR-25 family members and two anti-miRs that simultaneously targeted the two miR-19 family members. Finally, since the two miR-18 family members have a nearly identical sequence, we used one anti-miR that simultaneously inhibit both members.

*Pkd1*-KO mice were randomly assigned to receive either PBS, anti-miR-17, anti-miR-18, anti-miR-19, or anti-miR-25 mixtures. A dose of 20 mg/kg per injection was administered intraperitoneally at P10, P11, P12, and P15, and mice were sacrificed at P18. Each mouse injected with an anti-miR cocktail was internally controlled with at least one littermate receiving PBS, thus leading to a PBS control group with a larger sample size. Q-PCR analysis showed that anti-miR mixture targeting the miR-17 family reduced expression of all miR-17 family members but did not affect the expression of miR-18a, miR-19a, or miR-25 **(Fig. 2A, Supplementary Fig. 1A)**. Similarly, treatment with anti-miR-18, anti-miR-19, and anti-miR-25 mixtures specifically reduced the expression of miR-18, miR-19 and miR-25 family members, respectively, without affecting the expression of unrelated miRNAs **(Fig. 2B-D and Supplementary Fig. 1)**. To further rule out cross reactivity, we analyzed the expression of miR-21, an abundantly expressed, pathogenic miRNA in PKD.^8, 11, 14, 25^ miR-21 levels did not change in kidneys from mice treated with PBS compared to kidneys from mice treated with either anti-miR-17, 18, 19, or 25 mixtures **(Fig. 2E)**. A heat map summarizing these results is shown in **Fig. 2F**. Taken together, these results indicate that anti-miRs specifically target the cognate miRNA family members without affecting the expression of unrelated miRNAs.

**Figure 2:**
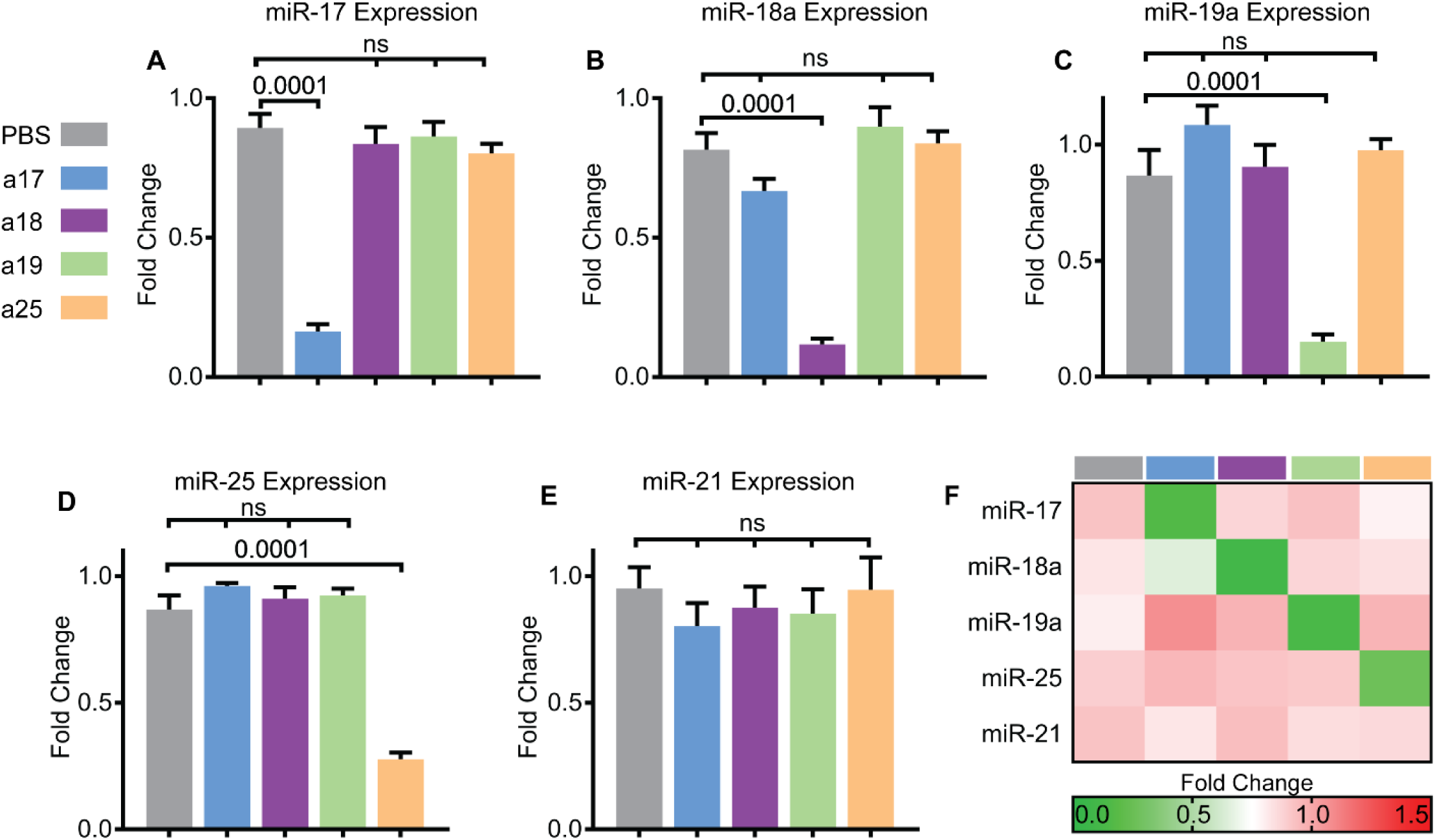
Anti-miR mixtures specifically inhibit cognate miRNAs. *Pkd1*-KO mice were randomized to receive PBS, anti-miR-17 (a17), anti-miR-18 (a18), anti-miR-19 (a19), or anti-miR-25 (a25) family inhibitors. **(A)** Q-PCR analysis revealed that the expression of miR-17 was reduced only in mice treated with anti-miR-17 but not in mice treated with anti-miRs against other families. **(B-D)** Similarly, miR-18, miR-19, and miR-25 levels were reduced only in mice treated with the respective anti-miRs. **(E)** Q-PCR analysis showed that anti-miR treatment did not affect the levels of miR-21, an abundantly expressed pathogenic miRNA in ADPKD, further demonstrating no off-target effects. **(F)** Heat map summary of Q-PCR data is shown. Data are presented as mean ± SEM, n=8 per group. Statistical analyses: One-way ANOVA (post hoc analysis: Dunnett’s multiple comparisons test), ns indicates *P*>0.05.

### Anti-miR-17, but not anti-miR-18, anti-miR-19 or anti-miR-25, attenuates cyst growth

To determine the effects of individually inhibiting each miRNA family on ADPKD progression, we evaluated cyst burden in mice treated with anti-miRs compared to PBS. Kidney histology was improved in anti-miR-17 treated mice, but not in mice treated with anti-miR-18, anti-miR-19, or anti-miR-25 **(Fig. 3A)**. No significant differences in body weight were found during any point in this study between treated and untreated mice **(Supplementary Fig. 2)**. However, kidney-weight-to-body-weight ratio (KW/BW) was reduced by 30.5% in mice treated with anti-miR-17 mixtures compared to PBS. In contrast, KW/BW ratio was not reduced in mice treated with anti-miR-18, anti-miR-19, or anti-miR-25 **(Fig. 3B)**. In fact, anti-miR-19 family treatment increased KW/BW by 31.4%. Moreover, cyst-index was reduced by 35.2% in anti-miR-17-treated mice compared to PBS-treated mice, but no change in cyst index was observed in mice treated with anti-miR-18, anti-miR-19, or anti-miR-25 mixtures **(Fig. 3C)**. H&E of all mice are shown in Supplementary Figure 3. To assess renal function, we measured blood urea nitrogen (BUN) levels which were reduced by 23% in mice treated with anti-miR-17 compared to PBS **(Fig. 4A),** however this observation did not reach statistical significance. No improvement in BUN was observed in mice treated with other anti-miRs mixtures. Phenotypic data of all mice analyzed in this study are shown in Supplementary table 2.

**Figure 3:**
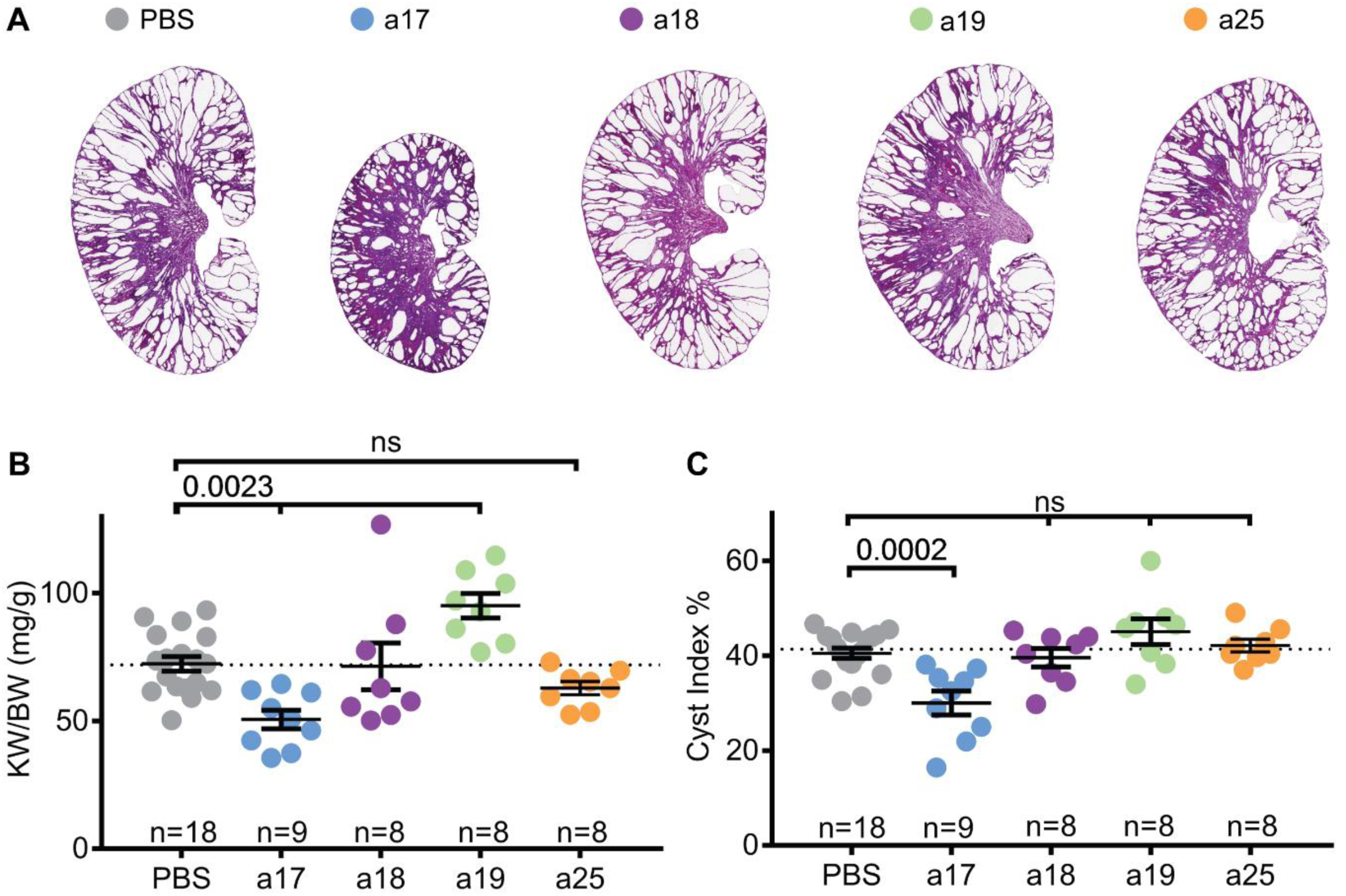
Anti-miR-17 but not anti-miR-18, anti-miR-19, or anti-miR-25 reduced cyst progression. **(A)** Representative H&E-stained kidney sections from *Pkd1*-KO mice treated anti-miR-17 (a17), anti-miR-18 (a18), anti-miR-19 (a19), or anti-miR-25 (a25) mice are shown. Only mice treated with anti-miR-17 showed reduction in kidney size. **(B)** Kidney-weight-to-body weight ratios (KW/BW) and **(C)** cyst index were reduced only in mice treated with anti-miR-17 treated mice compared to PBS. Data are presented as mean ± SEM. Statistical analyses: One-way ANOVA (post hoc analysis: Dunnett’s multiple comparisons test), ns indicates *P* >0.05.

**Figure 4:**
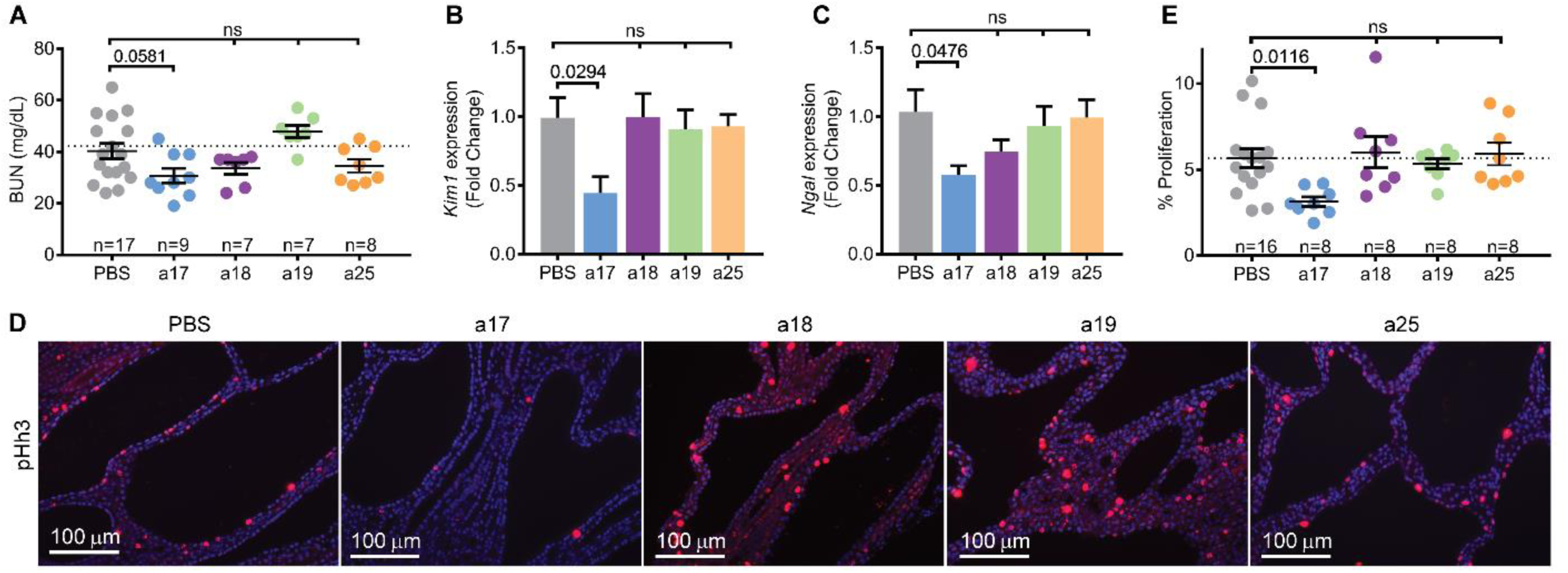
Anti-miR-17 improves renal function and reduces kidney injury and cyst proliferation. (**A)** Blood urea nitrogen (BUN) was decreased by 23% in anti-miR-17 treated mice compared to PBS treated mice. However this difference was not statically significant. (**B&C**) Q-PCR analysis revealed that expression of kidney injury markers, *Kim1* and *Ngal* were also reduced only in kidneys of anti-miR-17-treated mice. (N=6 per group) **(D&E)** To assess proliferation, kidney sections were stained using an antibody against phosphohistone-H3 (pHh3), a marker of proliferating cells. Quantification of PHh3 positive cells from ten random high powered images (20X) from each kidney section revealed that only anti-miR-17-treated mice showed a reduction in the number of proliferating cyst cells. Data are presented as mean ± SEM. Statistical analyses: One-way ANOVA (post hoc analysis: Dunnett’s multiple comparisons test), ns indicates *P* >0.05.

Anti-miR-17 also reduced kidney injury assessed by measuring the expression of kidney injury markers, *Kim1* and *Ngal*. Q-PCR analysis revealed a 55.1% reduction in *Kim1* and a 44.1% reduction in *Ngal* only in anti-miR-17 treated mice **(Fig. 4B, C)**. Next, we determined whether anti-miR-17 affected cyst proliferation. The number of cyst epithelial cells expressing phospho-histone H3, a marker of mitosis, was reduced by 44.6% in anti-miR-17 treated compared to PBS treated mice **(Fig. 4D, E)**. No change in cyst proliferation was observed in other groups. Thus, our results indicate that treatment with anti-miR-17, but not anti-miR-18, anti-miR-19, or anti-miR-25 mixture, reduced cyst progression and improved kidney function. These results suggest that within miR-17~92 and related clusters, the miR-17 family is the pathogenic element and the primary contributor to cyst progression.

### Anti-miR-17 treatment recapitulates the gene expression pattern observed after miR-17~92 deletion in *Pkd1*-KO kidneys

To understand the mechanism by which anti-miR-17 mediates its cyst-reducing effects, we began by performing RNA-seq analysis to compare gene expression profiles between kidneys of anti-miR-17 treated mice and PBS treated mice (n=3, each group). Pathway analysis of the differentially expressed genes revealed that the primary consequence of anti-miR-17 treatment was upregulation of mitochondrial metabolism pathways (oxidative phosphorylation (OXPHOS), mitochondrial function, fatty acid oxidation (FAO)) and downregulation of the inflammation and fibrosis pathways (atherosclerosis signaling, granulocyte adhesion, acute response signaling, etc.) **(Fig. 5A)**. Ingenuity pathway analysis software was used to identify the upstream regulators (URs) that could be responsible for the gene expression changes observed after anti-miR-17 treatment. This analysis showed that mitochondrial and metabolism-related gene networks regulated *Ppara, Pparg, Ppard,* and others were activated. In contrast, inflammation-associated gene networks were inhibited including those regulated by *Il1b* and *Tnf*. We have previously performed RNA-Seq analysis and identified genes that are differentially expressed as a result of miR-17~92 genetic deletion in *Pkd1*-KO kidney.^15^ We intersected the two RNA-seq datasets to discover a common gene signature between the genetic (miR-17~92 deletion) and pharmaceutical (anti-miR-17 treatment) approaches. There were 292 common differentially expressed genes found in both datasets. Unbiased One-minus Pearson hierarchal clustering using these 292 genes segregated the samples into two groups. *Pkd1*-KO kidneys without miR-17~92 deletion and PBS-treated *Pkd1*-KO kidneys clustered as one group whereas *Pkd1*-miR-17~92 double knockout kidneys and anti-miR-17 treated *Pkd1*-KO kidneys segregated as the second group. Pathway analysis using the 292 common genes also showed activation of mitochondrial function/metabolism and inhibition of inflammation **(Fig. 5E)**. Moreover URs *Ppara* and *Pparg* were predicted to be activated whereas inflammation-associated gene networks regulated by *Il1b, Tnf,* and other were predicted to be inhibited. Collectively, these results indicate that inhibition of the miR-17 family largely recapitulates the gene expression patterns observed due to miR-17~92 deletion in *Pkd1*-KO kidneys.

**Figure 5:**
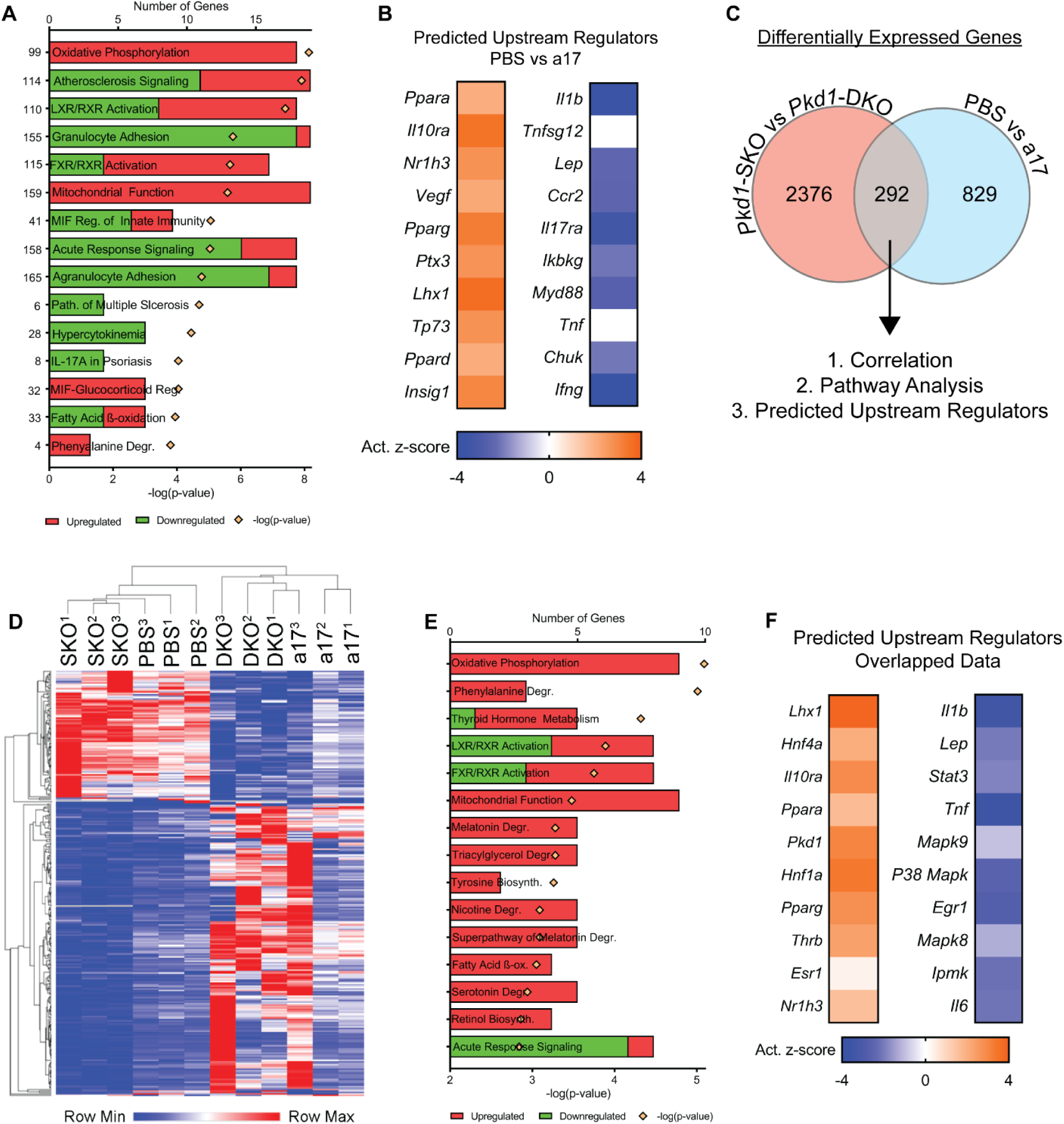
Anti-miR-17 treatment recapitulates the gene expression pattern observed after miR-17~92 deletion. RNA-seq analysis was performed using kidney RNA from PBS-treated and anti-miR-17-treated mice (a17) (n=3). **(A)** The top differentially regulated pathways in anti-miR-17-treated mice compared to PBS-treated mice are shown. Red indicates activation and green indicates inhibition of the indicated pathways **(B)** Ingenuity pathway analysis software was used to identify upstream regulators (URs) that may underlie the changes observed in gene expression. The activation z-scores of the top 10 most significantly upregulated (orange) and downregulated URs (blue) are shown. Positive z-scores (orange) indicate activation whereas negative z-scores (blue) indicate inhibition. **(C)** To determine if there is a common gene signature underlying the reduction in cyst growth observed between the anti-miR-17 and miR-17~92 genetic deletion, previous RNA-seq data comparing *Pkd1*^*F/RC*^ (*Pkd1*-SKO) and *Pkd1*^*F/RC*^; miR-17~92-KO (*Pkd1*-DKO) was intersected with the current data. There were 292 common, differentially regulated genes between the two data sets. **(D)** Unbiased hierarchical clustering of these 292 genes is shown. *Pkd1*-SKO and PBS-treated *Pkd1*-KO kidneys clustered as one group whereas *Pkd1*-DKO and anti-miR-17-treated *Pkd1*-KO kidney clustered together as a second independent group. **(E&F)** Pathways and UR analysis of the common gene signature also revealed an increase in mitochondrial metabolism and reduction in inflammation.

### Anti-miR-17 normalized kidney metabolism and reduced inflammation

Based on the unbiased analysis of our RNA-Seq data, we reasoned that anti-miR-17 treatment attenuates cyst growth by improving cyst metabolism and inhibiting cyst-associated inflammation. Q-PCR analysis revealed that expression of miR-17 targets *Ppara* and *Ppargc1a* was increased by 38.6% and 30.7%, respectively, in anti-miR-17-treated kidneys compared to PBS-treated kidneys **(Fig. 6A)**. PPARA and PPARGC1A are transcription factors that promote the expression of a large network of mitochondrial metabolism-related genes.^26-29^ Similarly, anti-miR-17 treatment also increased the expression of gluconeogenesis genes *Fbp1* and *G6pc2* by 62.4% and 70%, respectively **(Fig. 6A)**. To determine if genes encoding proteins for the electron transport chain (ETC) were increased, we analyzed expression of genes encoding subunits of each complex in the ETC **(Fig. 6B)**. *Ndufv1* (NADH dehydrogenase flavoprotein 1) and *Ndufa2* (NADH dehydrogenase 1 alpha subcomplex subunit 2) are both found in complex I^30, 31^ and were upregulated by 90.1% and 209.2% respectively. *Ppara* target genes *Etfa* (Electron Transfer Flavoprotein Alpha) and *Etfdh* (Electron Transfer Flavoprotein Dehydrogenase) are found in complex II^32^ and were upregulated by 124.3% and 26.8% respectively. *Cox5a* (Cytochrome c oxidase subunit 5a) found in complex IV^33^ was increased by 347.8%. *Atp5e* encodes a subunit of ATP synthase in complex V^34^ and was increased by 854.1%. Moreover, Western blot analysis confirmed that a critical subunit of ATP synthase, ATP5A was increased in anti-miR-17 treated mice compared to PBS treated mice **(Fig. 6C)**. To further characterize the metabolic phenotype of anti-miR-17 treated mice, we analyzed proteins in the mTOR pathway. This pathway is well known to regulate both glycolysis and mitochondrial metabolism^35-37^, and could partially explain the improvement of metabolism in anti-miR-17 treated mice. Western blot analysis revealed that both the active and total mTOR **(Fig. 6C)** levels were reduced in anti-miR-17 treated mice. Moreover, the expression of total and phosphorylated forms of 4EBP1 and S6RP, as well as P70S6K protein expression, were significantly decreased in anti-miR-17 treated kidneys compared to PBS treated kidneys. Defective autophagy is observed in ADPKD models.^38^ Inhibition of the mTOR pathway and improving cyst-metabolism has been shown to induce autophagy.^39^ Consistent with these observations, western blot analysis revealed a downregulation of P62 and upregulation of both LC3-I and LC3-II, suggesting an increase in autophagy in anti-miR-17-treated kidneys compared to PBS-treated kidneys.

**Figure 6:**
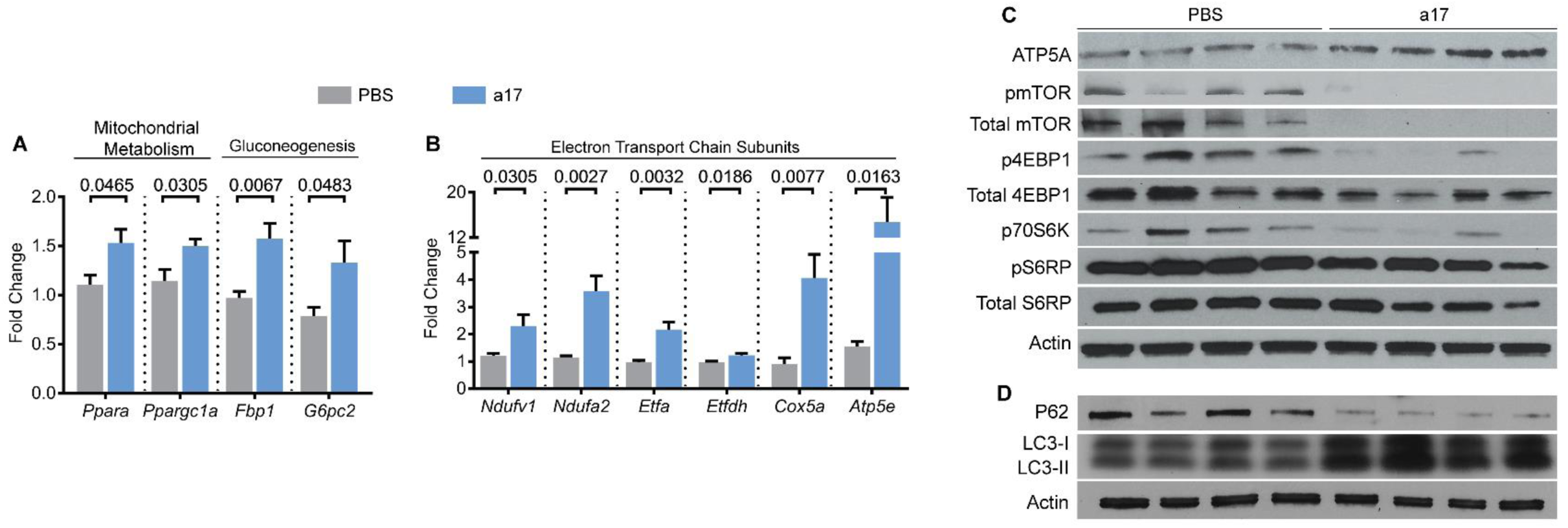
Anti-miR-17 upregulated metabolism-related genes, suppressed mTOR, and increased autophagy pathways. **(A&B)** Q-PCR analysis revealed that the expression of genes involved in various aspects of mitochondrial metabolism are increased in anti-miR-17-treated kidneys compared to PBS-treated kidneys. **(C)** Western Blot analysis showed an increase in ATP5A, a protein subunit of ATP synthase in anti-miR-17 treated mice compared to PBS-treated mice, further demonstrating an improvement in mitochondrial function. Additionally, mTOR and its downstream targets were significantly decreased. **(D)** Western Blot analysis of P62, LC3-I, and LC3-II revealed and increase in cellular autophagy in anti-miR-17 kidneys compared to PBS-treated kidneys. Data are presented as mean ± SEM. N=5 per group. Statistical analyses: Student’s t-test, ns indicates *P* >0.05.

Based on the RNA-Seq data, next, we validated whether the inflammatory pathways were downregulated in anti-miR-17-treated mice. Q-PCR analysis revealed that the expression of *Acta2* and *Col1a1*, two markers of fibrosis was decreased by 23.8% and 34.9%, respectively, in anti-miR-17 treated mice compared to PBS treated mice **(Fig. 7A)**. Furthermore, expression of cytokines *Tgfb2, Ifng, Ccl5, Ccl22, Il6,* and *Mip2* was decreased by 34%, 30.8%, 38.7%, 47.5%, 40.8%, and 36.5%, respectively **(Fig. 7A)**. Macrophages surrounding cysts consist of a heterozygous population of M1-like and M2-like macrophages, with the latter of the two thought to promote cyst growth^40, 41^. Q-PCR analysis showed that the markers of the pathogenic M2-like macrophages, *Arg1* and *Mrc1*, were decreased by 67.2%%, and 34.9%. Moreover, western blot analysis also revealed reduced expression MRC1 **(Fig. 7B)**. Lastly, immunohistochemistry analysis showed that the number of MRC1-positive macrophages was reduced by 54.1% in anti-miR-17 treated kidneys compared to PBS treated kidneys **(Fig. 6C&D)**. Thus, anti-miR-17 mediates its therapeutic effects by increasing mitochondrial metabolism through *Ppara*, reducing mTOR signaling, increasing autophagy, and decreasing inflammatory response notably through reduction of M2-like macrophage polarization.

**Figure 7:**
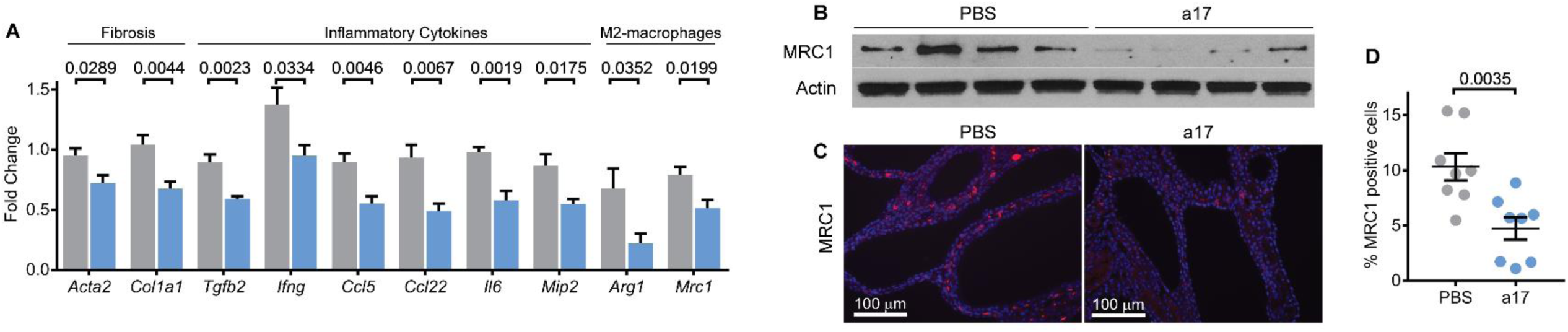
Anti-miR-17 treatment reduced fibrosis, inflammation, and M2-like macrophage. Q-PCR analysis demonstrated that the expression of inflammation and fibrosis-related genes was decreased in kidneys of anti-miR-17 treated mice. (N=5 per group) **(B)** Western blot analysis revealed that MRC1, a marker of M2-like macrophages, was decreased in anti-miR-17-treated kidneys compared to PBS-treated kidneys. **(C&D)** To further assess cyst-associated inflammation, kidney sections from anti-miR-17 and PBS-treated mice were stained using an antibody against MRC1. Quantification revealed MRC1-positive inflammatory cells were reduced anti-miR-17 treated compared to PBS-treated kidneys. (N=8 per group). Data are presented as mean ± SEM. Statistical analyses: Student’s t-test, ns indicates *P* >0.05.

## Discussion

We have previously shown that genetic inactivation of the miR-17~92 cluster attenuates disease progression in multiple ADPKD animal models.^15^ Thus, this cluster has emerged as a novel therapeutic target for ADPKD. However, the genetic approach reduced the expression of all miRNAs encoded by miR-17~92 precluding analysis of the individual pathogenic contributions of these miRNAs. Therefore, miRNAs within this cluster that should be therapeutically targeted have thus far remained unknown. We performed an *in-vivo* anti-miR screen to individually inhibit either miR-17, miR-18, miR-19, or miR-25 families in an orthologous mouse model of ADPKD. Treatment with anti-miR-17 family inhibitors recapitulated the anti-proliferative, cyst-reducing effects of deleting the entire miR-17~92 cluster. In contrast, inhibiting miR-18, miR-19, or miR-25 families had no impact on cyst growth. Hence, our results argue against functional cooperation between the various families in promoting cyst growth and instead point to the miR-17 family as the primary driver of PKD pathogenesis. These results suggest that rather than a complicated drug discovery process to simultaneously inhibit different families within the miR-17~92 cluster, a more simplified approach focused on targeting the miR-17 family alone will be sufficient in ADPKD. Our results provide a clear stepping stone for developing pharmaceutical grade anti-miRs against the miR-17 family.

miR-17~92 gene produces a single transcript, which gives rise to six individual mature miRNAs.^23^ However, the six miRNAs are processed with varying efficiencies producing different amounts of mature miRNAs.^42, 43^ Consistent with this notion, even though miR-17~92 gene is transcriptionally activated in ADPKD, the eventual level of mature miR-17 upregulation is much more pronounced than other miRNAs derived from this transcript. In fact, the miR-17 family alone accounts for more than 2.7% to the total dysregulated pool of all miRNAs, not just those produced by miR-17~92.^15^ In contrast, other miR-17~92 cluster families have minimal or no impact to the dysregulated miRNA pool in ADPKD models. This outsized contribution to the pathogenic miRNA pool may provide one explanation for why the miR-17 family appears to be the sole driver of cyst growth within the miR-17~92 cluster. Similar observations of specialization among members of this cluster have been made in other studies. Feingold syndrome, a genetic disease primarily characterized by short stature and skeletal abnormalities, can be caused due to germline microdeletions involving miR-17~92.^44^ The deletion of miR-17 seed family is sufficient to reproduce the skeletal malformations observed in Feingold syndrome, whereas individually deleting the other seed families has no effect. Similarly, the miR-17~92 cluster is essential for early B-cell development, and miR17/18 seed families appear to be solely responsible for this effect.^45^ Finally miR-17~92 cluster is amplified and oncogenic in lymphoma, and miR-19 overexpression is sufficient to fully reproduce this oncogenicity.^46^ Interestingly, despite this well-described role of miR-19 in promoting proliferation and cancer growth, we found that in the context of ADPKD, it does not have a significant pathogenic affect.

The second major insight from our work is that anti-miR-17 treatment also recapitulated the mechanistic effects of miR-17~92 genetic deletion. We identified a common gene signature between anti-miR-17 and miR-17~92 genetic deletion in ADPKD models. Importantly, consistent with our earlier observations, the primary cellular consequence of anti-miR-17 treatment was improved expression of metabolism-related gene networks, including the upregulation of direct miR-17 target *Pparα.* Thus, one of the direct mechanisms by which anti-miR-17 mediates its cyst-reducing effects may be through improvement in cyst metabolism. We have recently shown that PPARA upregulation is sufficient to improve cyst metabolism and attenuate cyst growth.^26^ Hence, stabilizing *Pparα* mRNA transcript could be a complementary or an alternative therapeutic approach for ADPKD. Our results also suggest that non-invasive assessment of various metabolic parameters could serve as potential pharmacodynamic biomarkers that predict anti-miR-17 function. Another consequence of anti-miR-17 treatment or miR-17~92 genetic deletion is the inhibition of various inflammation-related gene networks. We extended this observation and found that anti-miR-17 reduced cyst-associated macrophages particularly the M2-like macrophages, which have been shown to promote cyst growth.^40^ Whether the inhibition of inflammation is a direct or indirect effect of anti-miR-17 treatment remains unknown. As new mechanistic insights, we demonstrate that anti-miR-17 treatment also inhibits the mTOR signaling and induces autophagy, two pathways that intimately linked to the metabolic state of the cell.

Several questions have remained unanswered. The purpose of this screen was to uncover the pathogenic components within the miR-17~92 cluster in the context of ADPKD. Therefore, we elected to perform a short-term (five day), in-vivo screen directed at miR-17~92 and related clusters. Whether sustained inhibition of miR-17 family would produce a long-term beneficial effect was not addressed. Additionally, whether sustained miR-17 inhibition is safe and efficacious over the long term was also not studied. In our previous study, we genetically deleted the miR-17~92 cluster in the *Pkd1*-KO mice. miR-17~92 deletion essentially restored normal life span in *Pkd1*-KO mice. Moreover, in that study long-term (~6-months) treatment of *Nphp3*^*pcy/pcy*^ mice, a slow growth cystic disease model, with an anti-miR-17 compound also demonstrated a sustained benefit in attenuating cyst growth. Thus, our work suggests that sustained miR-17 inhibition is safe and efficacious. A second limitation is that we performed this study in only one model of ADPKD. We chose the *Pkd1*-KO mouse model because the majority patients with ADPKD have a mutation in the *PKD1* gene. Furthermore, anti-miR treatment has never been performed in a *Pkd1*-KO model. Nevertheless, we have previously shown that genetic knockout of miR-17~92 or pharmaceutical inhibition of miR-17 both reduce cyst burden in orthologous *Pkd2*-KO mouse model. Therefore, miR-17 family is also likely to be the primary driver of PKD pathogenesis in *Pkd2*-KO mice. Lastly, we did not study synergism between the various miR-17~92 cluster families. These miRNA families have many overlapping mRNA targets. Perhaps inhibition of one miRNA family is not sufficient to slow cyst growth, but co-inhibition of two or more miRNA families could have an additive effect. However, our results argue against this possibility as genetic deletion of miR-17~92 and pharmaceutical inhibition with anti-miR-17 produced a similar gene signature.

In summary, our studies suggest that the pathogenic component of the miR-17~92 cluster lies within the miR-17 family. Moreover, inhibition of only the miR-17 family recapitulated the mechanistic effects of miR-17~92 genetic deletion. Thus, our study provides a strong rationale for developing drugs against the miR-17 seed in ADPKD.

## Materials and Methods

### Mice

KspCre/*Pkd1*^F/RC^ mice were used for these studies. At 10 days of age KspCre/*Pkd1*^F/RC^ mice were randomized to receive PBS, Anti-miR-17 family cocktail, Anti-miR-18 family cocktail, Anti-miR-19 family cocktail, and Anti-miR-25 family cocktail (20 mg/kg total). Based on our previous experience, power analysis (alpha <5% and power >80%) was performed *a priori* to determine the sample size (n of at least 8). Mice were given intraperitoneal injections at postnatal day (P) 10, 11, 12, and 15, and were sacrificed at P18. All experiments involving animals were approved by the UTSW Animal Care and Use Committee. All experiments were performed in accordance with Institutional Animal Care and Use Committee guidelines and regulations. Equal males and females were used in all groups for all studies.

### Anti-miRs

Anti-miRs were acquired from Exiqon (Denmark). *In vivo* grade locked nucleic acid-modified anti-miRs were used. Each anti-miR was dissolved in PBS to create a 2 mg/uL stock which was kept at −20 ° C. On the day of injection, each anti-miR was thawed to 4 ° C and added to its respective anti-miR cocktail. A total dose of 20 mg/kg for each cocktail was injected. See Supplementary Table 1 for individual inhibitor product number, sequence, and target information.

### Tissue harvesting and analysis

Mice were anesthetized under approved protocols, blood was obtained via cardiac puncture. The right kidney was flash frozen for molecular analysis, and the left kidney was perfused with cold PBS and 4% (wt/vol) paraformaldehyde and then harvested. Kidneys were fixed with 4% paraformaldehyde for 2 hours and then embedded in paraffin for sectioning. Sagittal sections of kidneys were stained with hematoxylin and eosin (H&E) for cyst index analysis using Image J software.

### Renal function tests

BUN was measured using the Vitros 250 Analyzer.

### RNA Isolation and Quantitative RT-PCR (Q-PCR)

Total RNA was isolated from mouse kidneys using miRNeasy Mini kits (Qiagen). First-strand cDNA was synthesized from mRNA using the iScript cDNA synthesis kit (Bio-Rad) and Superscript III (Invitrogen), and Q-PCR was performed using the iQ SYBR Green Supermix (Bio-Rad). The Universal cDNA Synthesis kit from Exiqon was used for first-strand synthesis from miRNA. Q-PCR was performed by using miRNA-specific forward and reverse locked nucleic acid (LNA)-enhanced PCR primers from Exiqon. The samples were loaded in duplicate on a CFX ConnectTM Real-time PCR detection system. 18S and SNORD68 RNA were used to normalize expression of mRNA and miRNA, respectively. Data were analyzed using the Bio-Rad CFX software. The sequences of the PCR primers are shown in Supplementary Table 2.

### RNA Sequencing

Total RNA was extracted from whole kidney lysate of three PBS treated and anti-miR-17 treated mice. Strand specific RNA-Seq libraries were prepared using the TruSeq Stranded Total RNA LT Sample Prep Kit from Illumina. After quality check and quantification, libraries were sequenced at the UT Southwestern McDermott Center using a Hiseq2500 sequencer to generate 51 bp single-end reads. Before mapping, reads were trimmed to remove low quality regions in the ends. Trimmed reads were mapped to the mouse genome (mm10) using TopHat v2.0.12 with the UCSC iGenomes GTF file from Illumina. Alignments with mapping quality less than 10 were discarded. Expression abundance estimation and differential expression gene identification was done using edgeR. Genes with a *P*-value < 0.05 were deemed significantly differentially expressed between the two conditions.

### Immunofluorescence Staining

The following antibodies and dilutions were used on paraffin embedded section for immunofluorescence staining: phosphohistone H3 (1:400, Sigma-Aldrich H0412), and anti-Mannose Receptor (Abcam ab64693, 1:400). Secondary antibodies were conjugated to Alexa Fluor 594 (Molecular Probes, 1:400). **Immunofluorescence Quantification**: Image J’s Find Maxima feature was utilized to determine the percentage of phospho-histone H3 or MRC1 positive cells from ten random high powered (20X) fields from each kidney section. An average of 6300 cells were counted per mouse. The total number of mice analyzed for phospho-histone H3 or MRC1 quantification are shown in figures 4 and 7, respectively.

### Western Blot analysis

Total protein was extracted from kidneys. 15 µg of protein was loaded on a 4-20% SDS-polyacrylamide gel and the proteins were transferred to a nitrocellulose membrane. The membrane was blocked with 5% milk and probed overnight at 4°C with antibodies (Dilution 1:1000): ATP5e (Abcam ab110413), pmTOR (Cell Signaling #2971), Total mTOR (Cell Signaling #2972), p4EBP1(Cell Signaling #2855), Total 4EBP1 (Cell Signaling #9644), pS6RP (Cell Signaling #2211), Total S6RP (Cell Signaling #2217), and Mannose Receptor 1 (Abcam ab64693). Goat-anti-rabbit HRP-conjugated IgG was used as the secondary antibody. The blot was developed using the SuperSignal West Dura Extended Duration substrate from Pierce. The protein bands were quantified using Quantity One imaging software from Bio-Rad.

### Clustering Analysis

Heat map and clustering analysis was performed using Morpheus software found at: https://software.broadinstitute.org/morpheus/

### Statistical Analysis

Statistical analysis was performed using One-way ANOVA. Next, a post hoc analysis (Dunnett’s one way multiple comparisons test) was performed using PBS as the control group. P-values indicate significant differences between PBS treatment and respective anti-miR treatment. Lastly, Student’s *t-* test was performed in analyses with two groups. Data are shown as the mean ± SEM.

## Author Contributions

M.Y., R.L, and V.P. conceived the idea. M.Y., R.L., and A.F. performed the experiments. M.Y. prepared the figures and wrote manuscript with V.P.

## Acknowledgments

We thank the UT Southwestern O’Brien Kidney Research Core Center (P30DK079328), the Mayo Translation PKD center (DK090728), UT Southwestern Bioinformatics core, Dr. Thomas Carroll (UT Southwestern), and Dr. Beth Levine (UT Southwestern) for providing critical reagents and services. We thank Dr. Silvia Ferrè (UT Southwestern) for helpful comments and discussion during the preparation of this manuscript. The work from the authors’ laboratory is supported by National Institute of Health grants R01DK102572 (to V.P).

## Supplementary Information

**Supplementary Figure 1:**
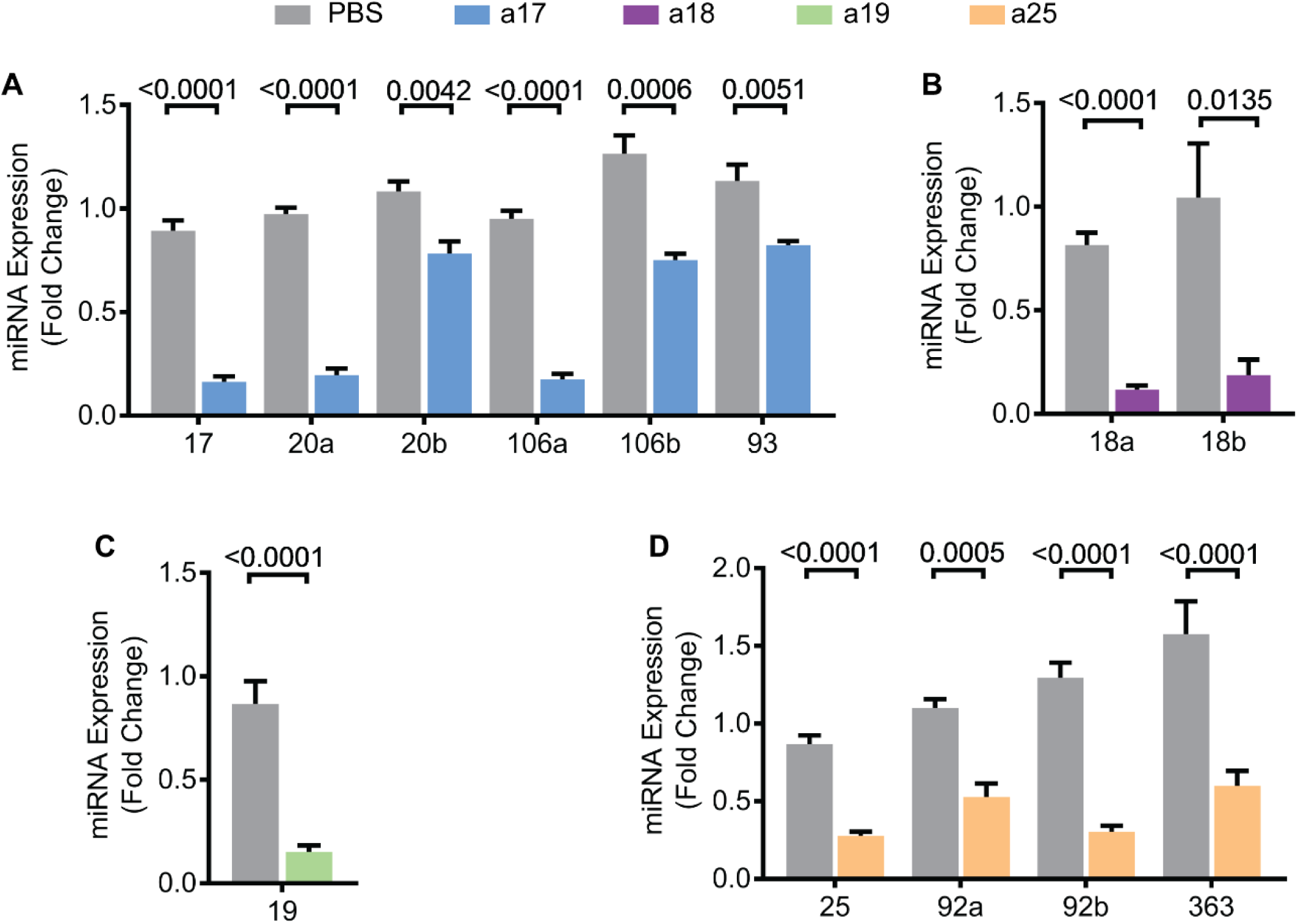
Anti-miRs significantly reduced expression of miRNAs in each target family. Q-PCR analysis was performed to determine the expression of each miRNA in the **(A)** miR-17, **(B)** miR-18, **(C)** miR-19, and **(D)** miR-25 families after inhibition with their respective anti-miR cocktails. Data are presented as mean ± SEM. N=5 per group. Statistical analyses: Student’s t-test, ns indicates *P* >0.05.

**Supplementary Figure 2:**
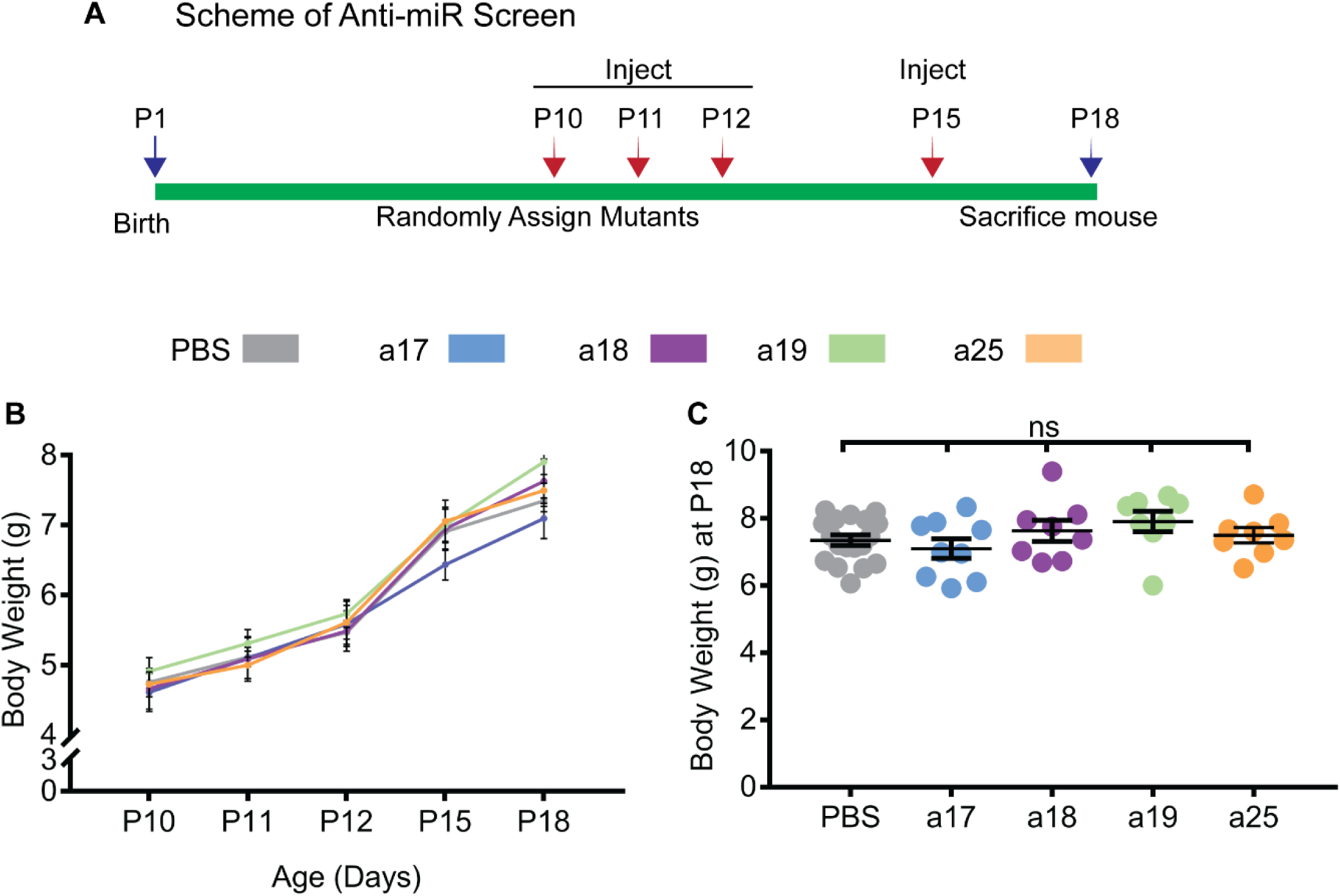
(A) Scheme of anti-miR screen in *Pkd1*-KO mice. Mice were randomly assigned to receive either PBS, anti-miR-17 (a17), anti-miR-18 (a18), anti-miR-19 (a19), or anti-miR-25 (a25). The total anti-miR dose per injection was 20 mg/kg. Each mouse was injected at P10, P11, P12, and P15, and sacrificed at P18. **(B)** Body weights were monitored during anti-miR treatment. There were no significant changes in body weights between treatment arms throughout the study. **(C)** Body weight of each mouse on P18 prior to sacrifice. There was no difference in body weight between groups. Statistican analysis: One-way ANOVA (post hoc analysis: Dunnett’s multiple comparisons test). ns indicates *P*>0.05.

**Supplementary Figure 3:**
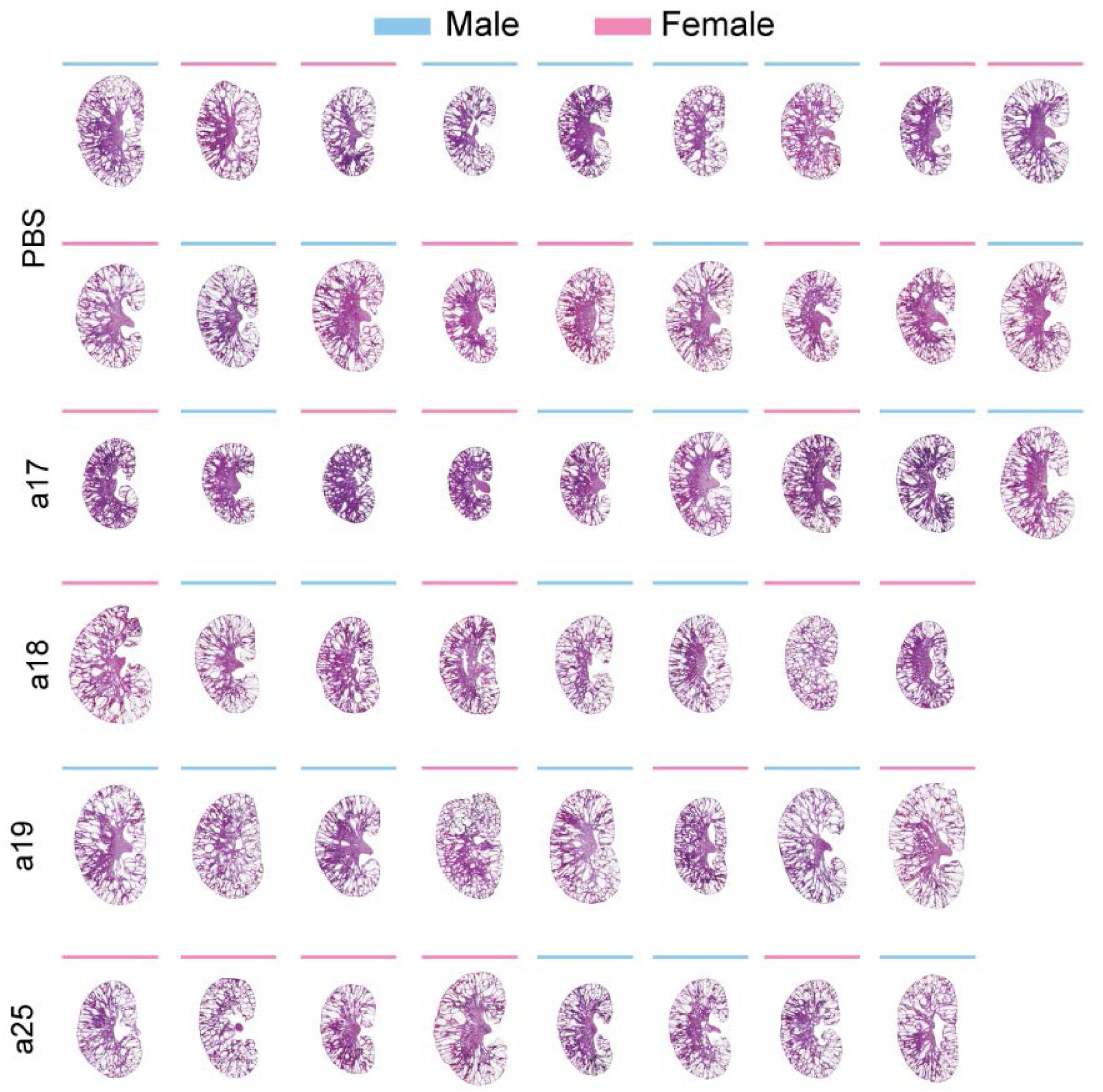
H&E kidney sections for all mice sacrificed in this study. PBS n=18, anti-miR-17 (a17) n=9, anti-miR-18 (a18) n=8, anti-miR-19 (a19) n=8, and anti-miR-25 (a25) n=8.

**Supplementary Table 1:**
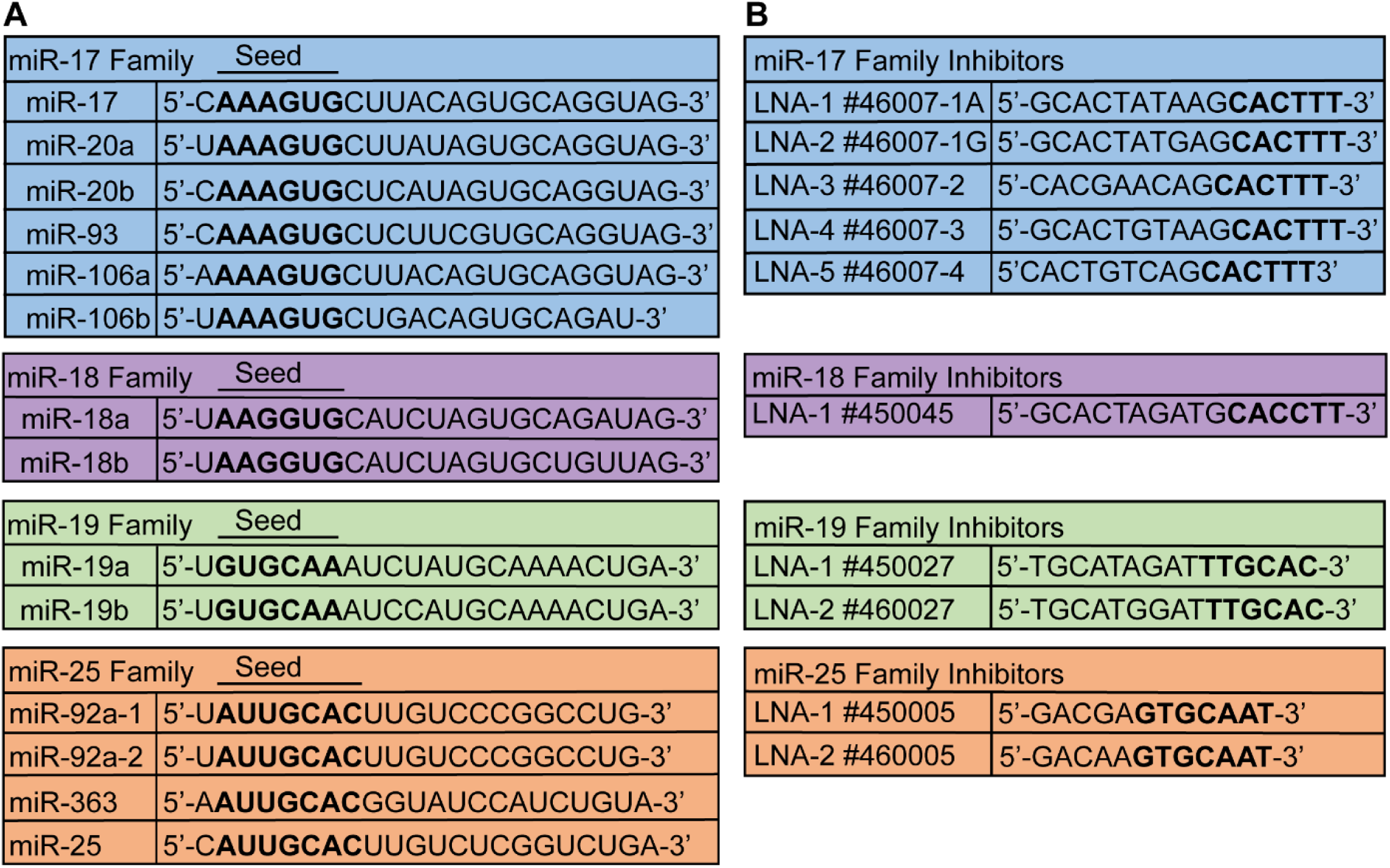
(A) Mature miRNA sequences of each family are shown. Bolded letters denote the seed sequence of each miRNA. **(B)** Sequences and product information of each inhibitor used in this study is shown. Bolded letters denote anti-miR sequence that is perfectly complimentary to the seed sequence of its corresponding miRNA family.

**Supplementary Table 2:** Full information of all mice used in this study

| PBS Treatment |  |  |  |  |
| --- | --- | --- | --- | --- |
| Sex | KW | KW/BW | Cyst Index | BUN |
| M | 527 | 76.35468 | 36.1% | 39 |
| F | 483 | 74.11386 | 43.3% | N/A |
| F | 383 | 65.03651 | 44.4% | 30 |
| M | 463 | 74.46124 | 43.9% | 35 |
| M | 338 | 62.75529 | 30.4% | 42 |
| M | 418 | 73.61747 | 46.7% | 54 |
| M | 516 | 83.73905 | 40.4% | 65 |
| F | 388 | 67.38451 | 34.9% | 48 |
| F | 612 | 90.74733 | 40.9% | 56 |
| F | 617 | 93.3293 | 45.5% | 48 |
| M | 395 | 61.62246 | 44.1% | 24 |
| M | 613 | 89.11179 | 44.9% | 55 |
| F | 326 | 50.30088 | 38.3% | 32 |
| F | 433 | 62.05216 | 43.6% | 34 |
| M | 586 | 82.94409 | 41.8% | 39 |
| F | 380 | 63.54515 | 38.8% | 25 |
| F | 395 | 59.13174 | 31.4% | 27 |
| M | 518 | 72.38681 | 40.0% | 32 |
| AVE | 466.17 | 72.37 | 40.5% | 40.29 |
| STDEV | 96.76 | 12.02 | 4.8% | 12.18 |

| Anti-miR-17 Treatment |  |  |  |  |
| --- | --- | --- | --- | --- |
| Sex | KW | KW/BW | Cyst Index | BUN |
| F | 242 | 37.42075 | 25.0% | 23 |
| M | 341 | 61.10016 | 28.9% | 30 |
| F | 296 | 46.26446 | 16.4% | 26 |
| F | 197 | 35.62387 | 21.9% | 28 |
| M | 282 | 50.91172 | 32.5% | 39 |
| M | 451 | 64.53921 | 38.1% | 39 |
| F | 386 | 55.09563 | 34.6% | 45 |
| M | 326 | 42.40375 | 35.3% | 19 |
| M | 423 | 62.16931 | 37.3% | 28 |
| AVE | 327.11 | 50.61 | 30.0% | 30.78 |
| STDEV | 83.20 | 10.86 | 7.5% | 8.48 |

| Anti-miR-18 Treatment |  |  |  |  |
| --- | --- | --- | --- | --- |
| Sex | KW | KW/BW | Cyst Index | BUN |
| F | 951 | 126.902 | 43.8% | N/A |
| M | 607 | 87.9455 | 40.4% | 37 |
| M | 377 | 52.4194 | 36.4% | 36 |
| F | 401 | 57.7145 | 34.5% | 37 |
| M | 495 | 77.5984 | 45.3% | 38 |
| M | 352 | 55.6522 | 42.4% | 26 |
| F | 376 | 62.8657 | 44.0% | 37 |
| F | 305 | 50.2058 | 29.8% | 24 |
| AVE | 483.00 | 71.41 | 39.6% | 33.57 |
| STDEV | 211.39 | 25.94 | 5.5% | 5.91 |

| Anti-miR-19 Treatment |  |  |  |  |
| --- | --- | --- | --- | --- |
| Sex | KW | KW/BW | Cyst Index | BUN |
| M | 665 | 92.8771 | 47.1% | 46 |
| M | 719 | 103.872 | 45.8% | 53 |
| M | 557 | 86.1829 | 38.3% | 47 |
| F | 705 | 109.049 | 46.5% | 57 |
| M | 732 | 114.806 | 48.1% | N/A |
| F | 400 | 76.9231 | 34.0% | 37 |
| M | 686 | 97.1121 | 60.0% | 46 |
| F | 600 | 80.3213 | 40.7% | 49 |
| AVE | 633.00 | 95.14 | 45.1% | 47.86 |
| STDEV | 111.72 | 13.61 | 7.8% | 6.28 |

**Supplementary Table 2:** Full information of all mice used in this study
| Anti-miR-25 Treatment |  |  |  |  |
| --- | --- | --- | --- | --- |
| Sex | KW | KW/BW | Cyst Index | BUN |
| F | 469 | 69.6673 | 45.6% | 45 |
| F | 380 | 53.5588 | 49.0% | 42 |
| F | 309 | 52.444 | 40.4% | 35 |
| F | 555 | 73.0167 | 42.9% | 41 |
| M | 372 | 59.6249 | 37.0% | 27 |
| M | 425 | 65.3846 | 40.4% | 30 |
| F | 408 | 62.8659 | 42.1% | 28 |
| M | 446 | 66.4878 | 39.7% | 29 |
| AVE | 420.50 | 62.88 | 42.1% | 34.63 |
| STDEV | 73.38 | 7.31 | 3.8% | 7.15 |

**Supplementary Table 3:**
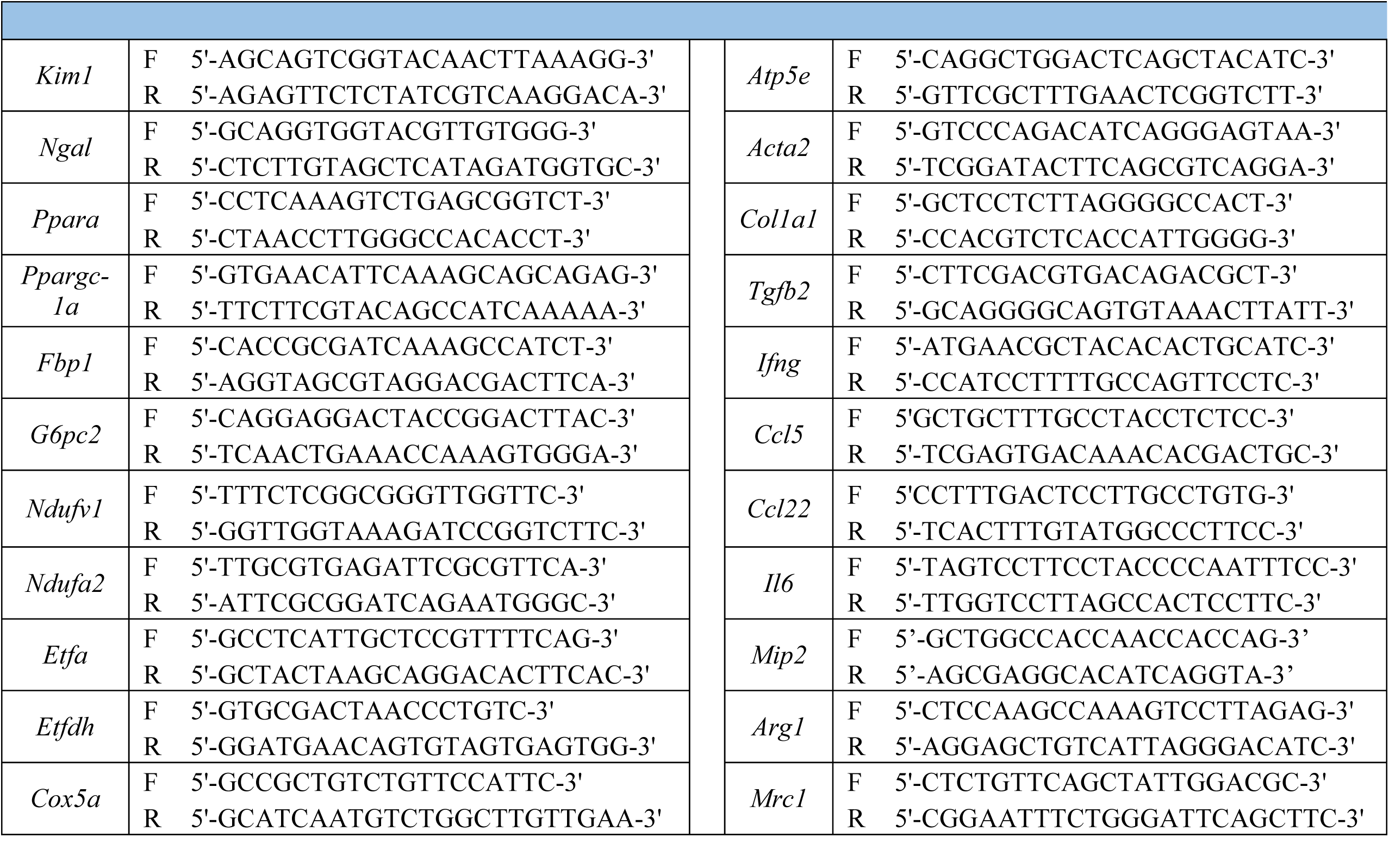
Sequences of primers used in the current study

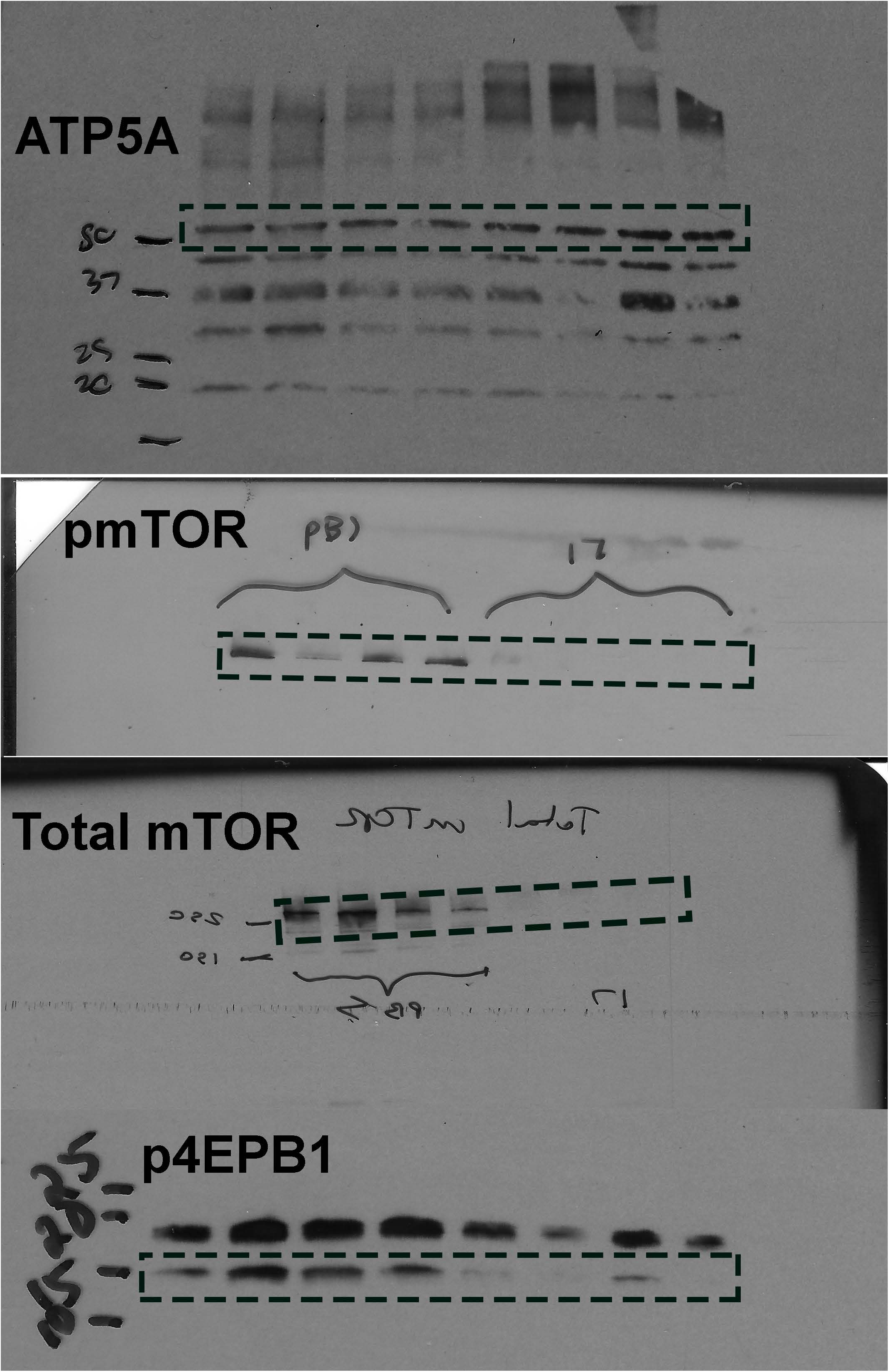

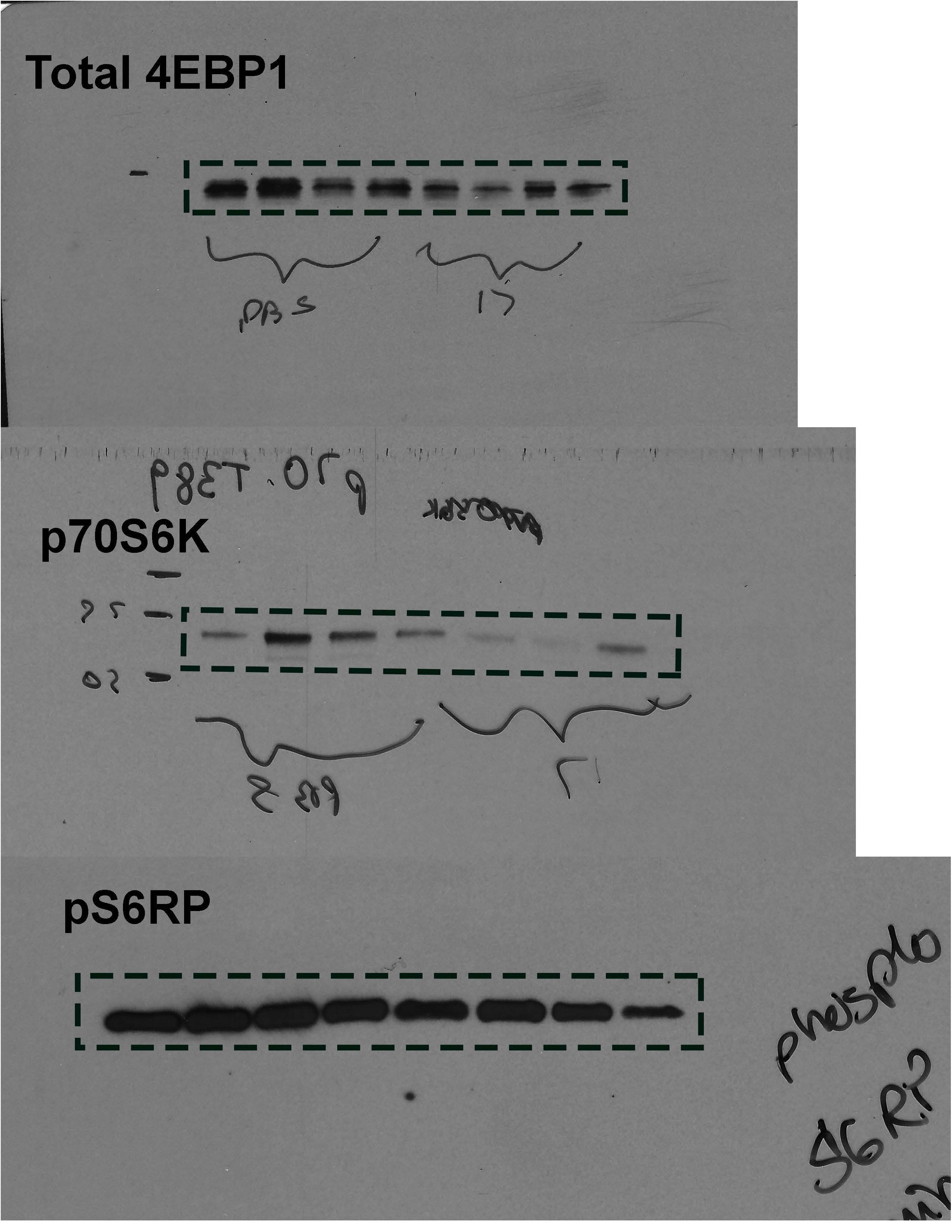

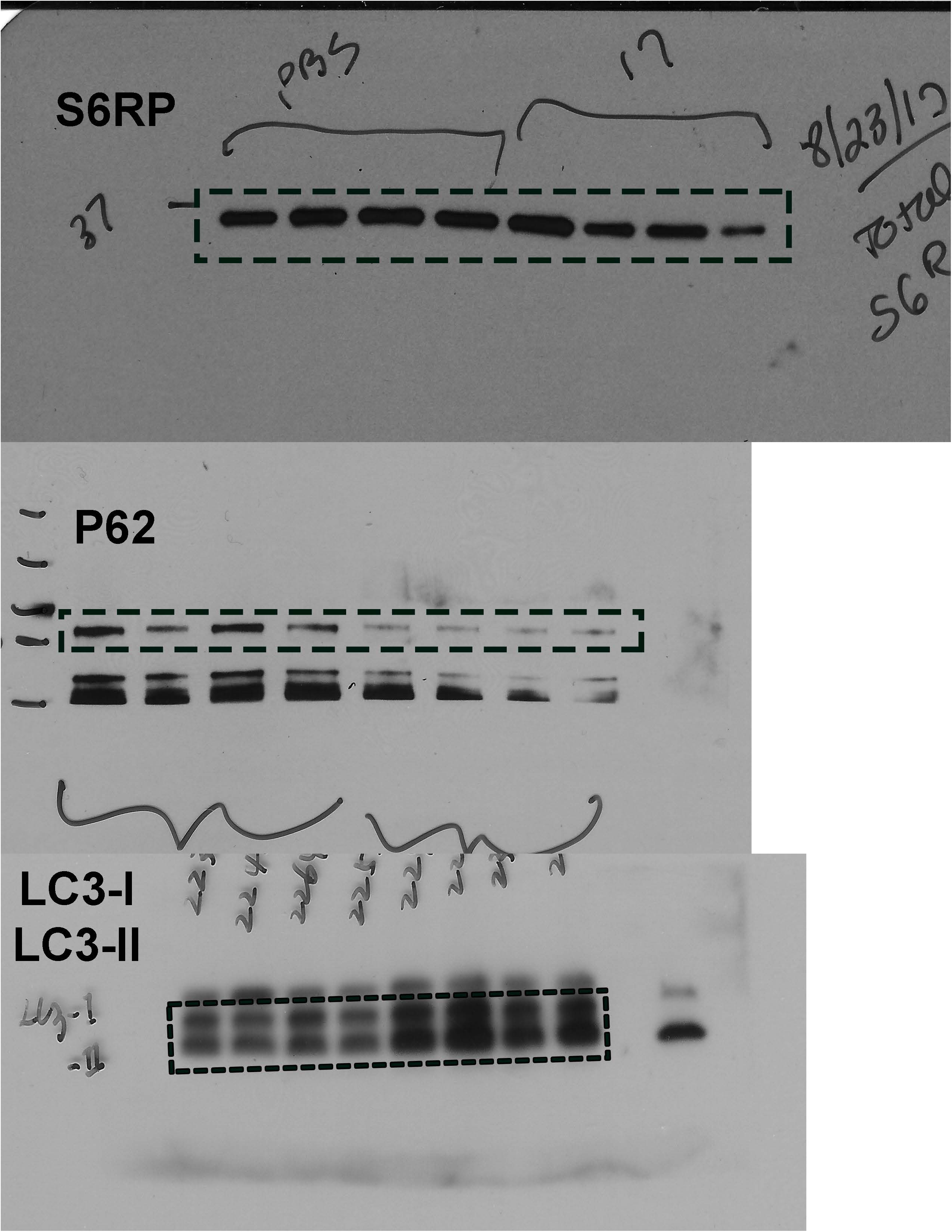

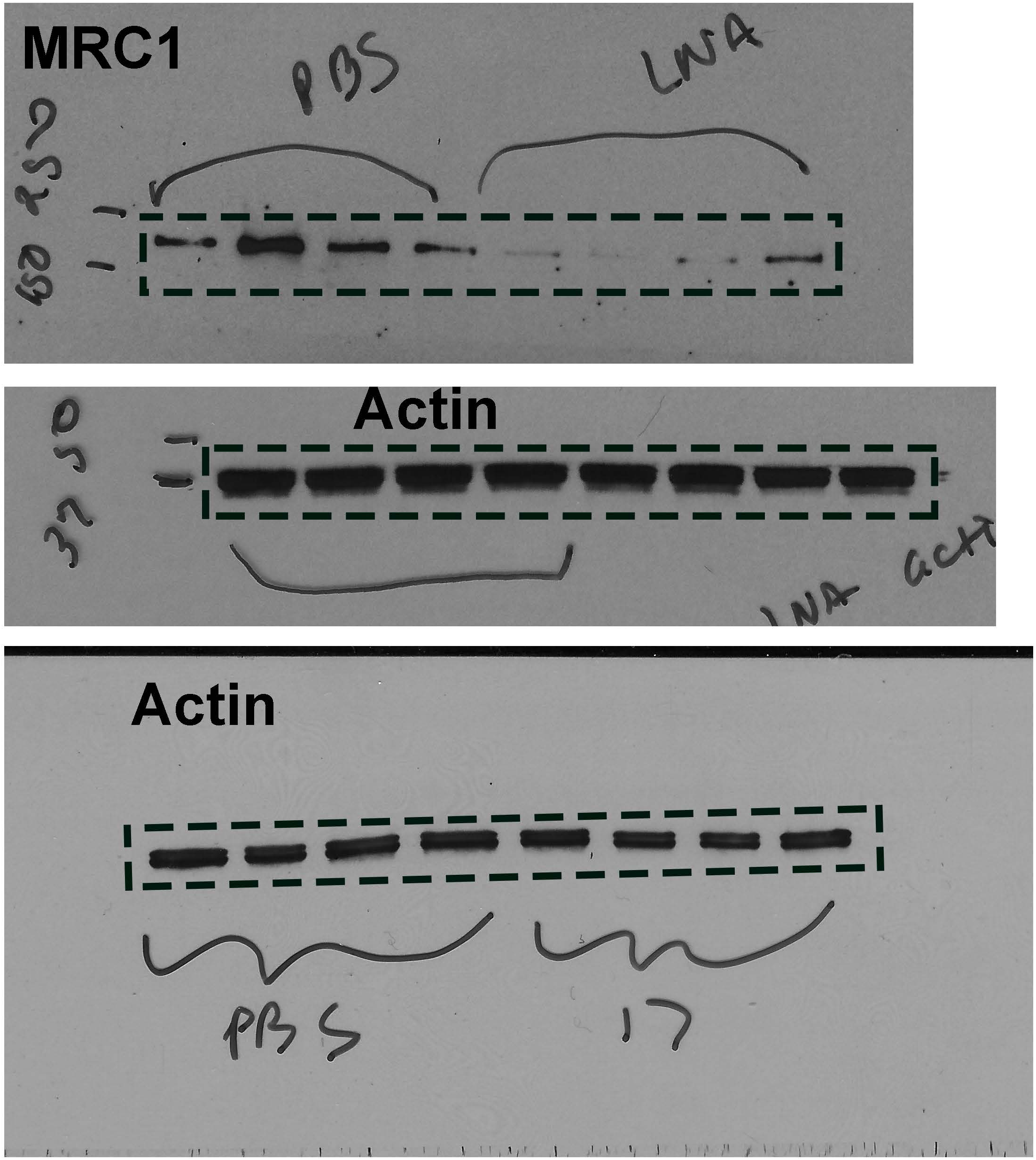

